# Functional insights into nucleoside diphosphate kinases encoded by two *ndk* paralogs in *Waddlia chondrophila*

**DOI:** 10.1101/2025.10.15.682642

**Authors:** Giti Ghazi-Soltani, Carole Kebbi-Beghdadi, Simone E. Adams, Gilbert Greub

## Abstract

The *Chlamydiota* phylum consists of obligate intracellular bacteria, including well-known pathogens and emerging environmental species, with diverse host ranges and metabolic capabilities. Among these bacteria, the gene, which encodes nucleoside diphosphate kinase (*ndk*), is present in variable copy numbers. While most chlamydial species carry a single copy of *ndk*, some species have two copies. In *W. chondrophila*, the two Ndk proteins encoded by *ndk* paralogs retain conserved kinase motifs but differ in subcellular localization, suggesting divergent functional roles. According to localization studies performed in heterologous expression systems, WcNdk1 is confined to the inclusion and probably supports nucleotide metabolism, while WcNdk2 localizes to the host nucleus, perinuclear space, and Golgi apparatus, suggesting involvement in host interaction. Azidothymidine (AZT), a known Ndk inhibitor, impaired *W. chondrophila* growth, potentially through inhibition of WcNdk2. However, the lack of genetic tools and the absence of *in vitro* enzymatic assays currently limit definitive functional conclusions. Our data suggest potential functions for Ndks in *W. chondrophila,* providing a foundation for future studies on Ndk-mediated interactions between this pathogen and its host.

## INTRODUCTION

The *Chlamydiota* phylum encompasses several families of obligate intracellular bacteria, including six main families: *Chlamydiaceae*, *Parachlamydiaceae*, *Waddliaceae*, *Simkaniaceae*, *Rhabdochlamydiaceae*, and *Criblamydiaceae*. Members of these families share a biphasic developmental cycle, which reflects their adaptation to intracellular environments and reliance on host cell machinery for survival.

*W. chondrophila* belongs to the family *Waddliaceae* and is known as an abortigenic agent in ruminants. It was first isolated from an aborted bovine fetus^1,2^ and serological studies showed a strong association between anti-*Waddlia* antibodies and bovine abortion^3^. In humans, *W. chondrophila* seropositivity is significantly associated with adverse pregnancy outcomes^4–6^. This pathogen was also detected in respiratory tract samples of patients with pneumonia and children with bronchiolitis^7,8^.

The *W. chondrophila* developmental cycle is biphasic and is divided into three major phases^9,10^. In the early phase (0-8 hours post-infection, hpi) elementary bodies (EBs) enter in host cells and differentiate into reticulate bodies (RBs). This phase is followed by a proliferation phase (8-24 hpi) during which the number of RBs increases exponentially through binary fission. In the late phase (24 - 48 hpi), RBs revert into EBs and lysis of the host cell releases infectious progeny. Under environmental or antibiotic stress, the developmental cycle is arrested leading to an enlarged, non-dividing RB-like form called Aberrant Body (AB)^11^. While *W. chondrophila* significantly impacts both animal and human health, the specific molecular and genomic pathways, particularly those involving developmental and regulatory proteins, remain largely unexplored. Functional characterization of proteins, such as nucleoside diphosphate kinase (Ndk), could help us to better understand the biology and pathogenicity of this pathogen and its adaptation inside host cells.

Ndk is a highly conserved enzyme that plays a central role in maintaining nucleotide pools within both bacterial and eukaryotic cells. The maintenance of intracellular NTP pools is vital for the survival and proliferation of all living organisms. Ndk regulates nucleotide pools through its autophosphorylation and phosphotransferase activities. This ensures that NTP and dNTP levels remain balanced for optimal cellular function.

Beyond its housekeeping functions, Ndk is implicated in various cellular processes, including cell differentiation and development in eukaryotes and prokaryotes^12–15^. Additionally, Ndk is involved in the regulation of gene expression^16–18^. Due to its ability to bind ATP, Ndk is involved in inhibiting extracellular ATP (eATP) and disrupting purinergic signaling in immune and inflammatory responses^19–21^.

We previously reported transcriptomic data for *W. chondrophila* comparing genes expression in RBs at 24 hpi and EBs at 72 hpi^22^. In that dataset, both *ndk* genes were differentially expressed between the two developmental stages, each showing a fold change of 0.3 in EBs compared with RBs. Despite the well-established functions of Ndk across various organisms, its specific function in the *Chlamydiota* remains uncharacterized. Based on these observations, we hypothesized that this enzyme could play an important role in the development and pathogenesis of *W. chondrophila*. In this study, we characterize the two Ndk proteins in this organism to better understand their potential role in the bacterium’s development. This study provides an overall view of Ndk proteins in the *Chlamydiota* phylum and offers functional insights into the Ndks proteins of *W. chondrophila*. Our results pave the way for more detailed research once genetic tools become available for *W. chondrophila*.

## RESULTS

### Evolution of the *ndk* operon in the *Chlamydiota* phylum

Similar to other organisms, all bacteria of the *Chlamydiota* phylum possess at least one copy of the *ndk* gene. A second copy (*ndk2*) is present in members of the *Parachlamydiaceae*, *Waddliaceae*, and *Criblamydiaceae* families (Figure 1A). Interestingly, *ndk2* forms an operon with the ancestral *ndk1* gene. In most species, this operon is further associated with *pabA*, which encodes para-aminobenzoate synthase and is involved in the folate biosynthesis pathway. Comparison of gene-based and species-based phylogenetic trees (Figure 1B, C) suggests that the duplication of *ndk2* most likely originated from a single gene duplication event in the common ancestor of the *Parachlamydiaceae*, *Waddliaceae*, and *Criblamydiaceae*. However, a scenario of ancestral duplication with subsequent loss in *Simkaniaceae* and *Chlamydiaceae* cannot be excluded, although it is less parsimonious.

**Figure1:**
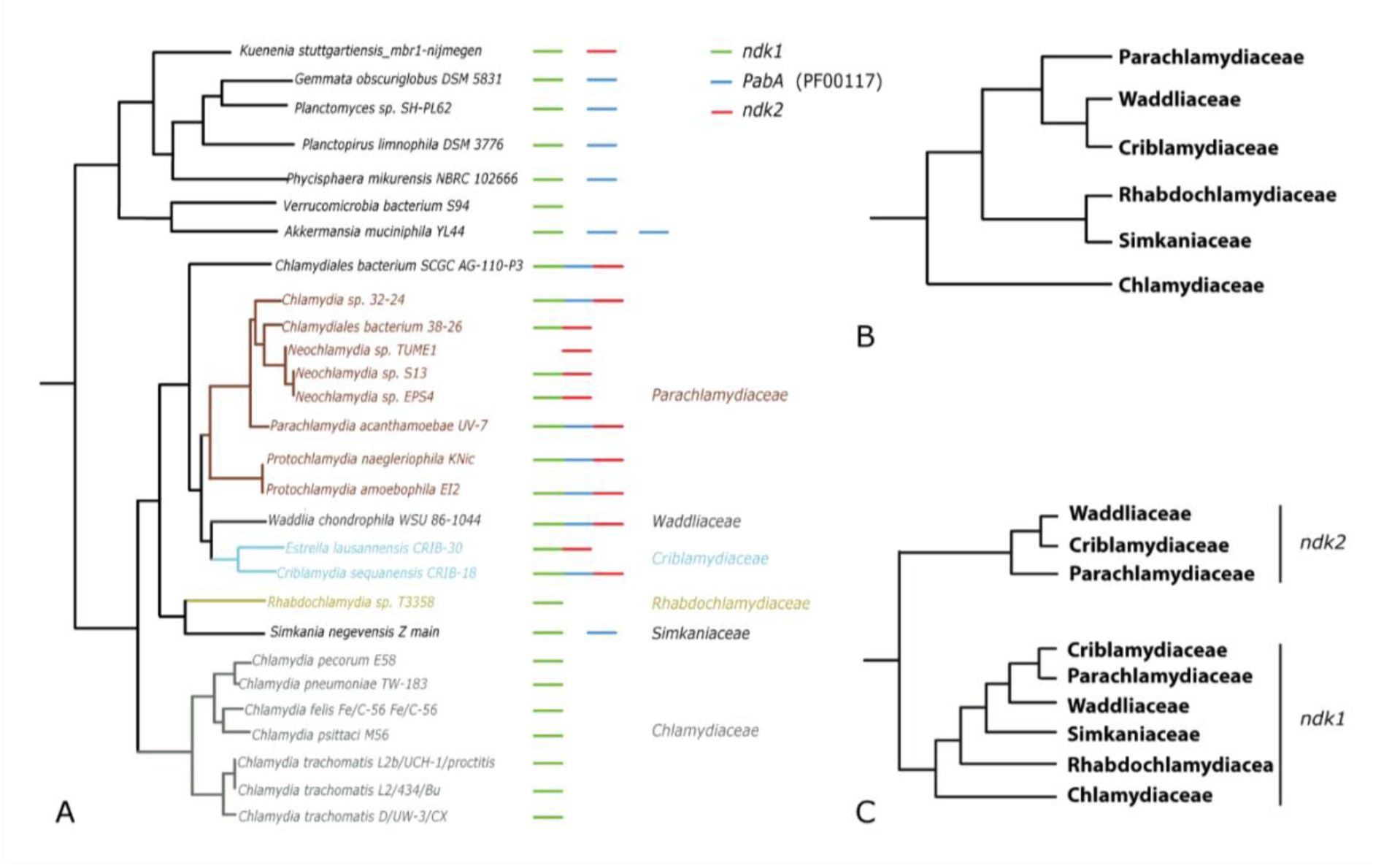
Phylogeny of *Chlamydiota* phylum and the *ndk* gene operon evolution. (A) Species phylogeny based on 32 single-copy orthologous genes. (B) Simplified representation of the species phylogeny highlighting major clades. (C) Phylogeny based on the *ndk* gene, showing its evolutionary relationship among *Chlamydial* species.

### High sequence and structural conservation of Ndk across species

The multiple sequence alignment shows a high degree of sequence conservation of Ndk proteins from various bacterial and eukaryotic species (Figure 2A). Specifically, the His-Gly-Ser-Asp (HGSD) motif, which is the enzyme’s active site^23^, is well conserved across all aligned sequences. The highly conserved histidine residue inside this motif transiently receives the phosphate group during autophosphorylation reaction and is, therefore, critical for the catalytic function of Ndk. In addition to the HGSD motif, other conserved residues are observed in regions associated with nucleotide binding and structural stability, many of which are enriched in hydrophobic amino acids. These residues likely contribute to the proper folding and enzymatic efficiency of Ndk.

**Figure 2:**
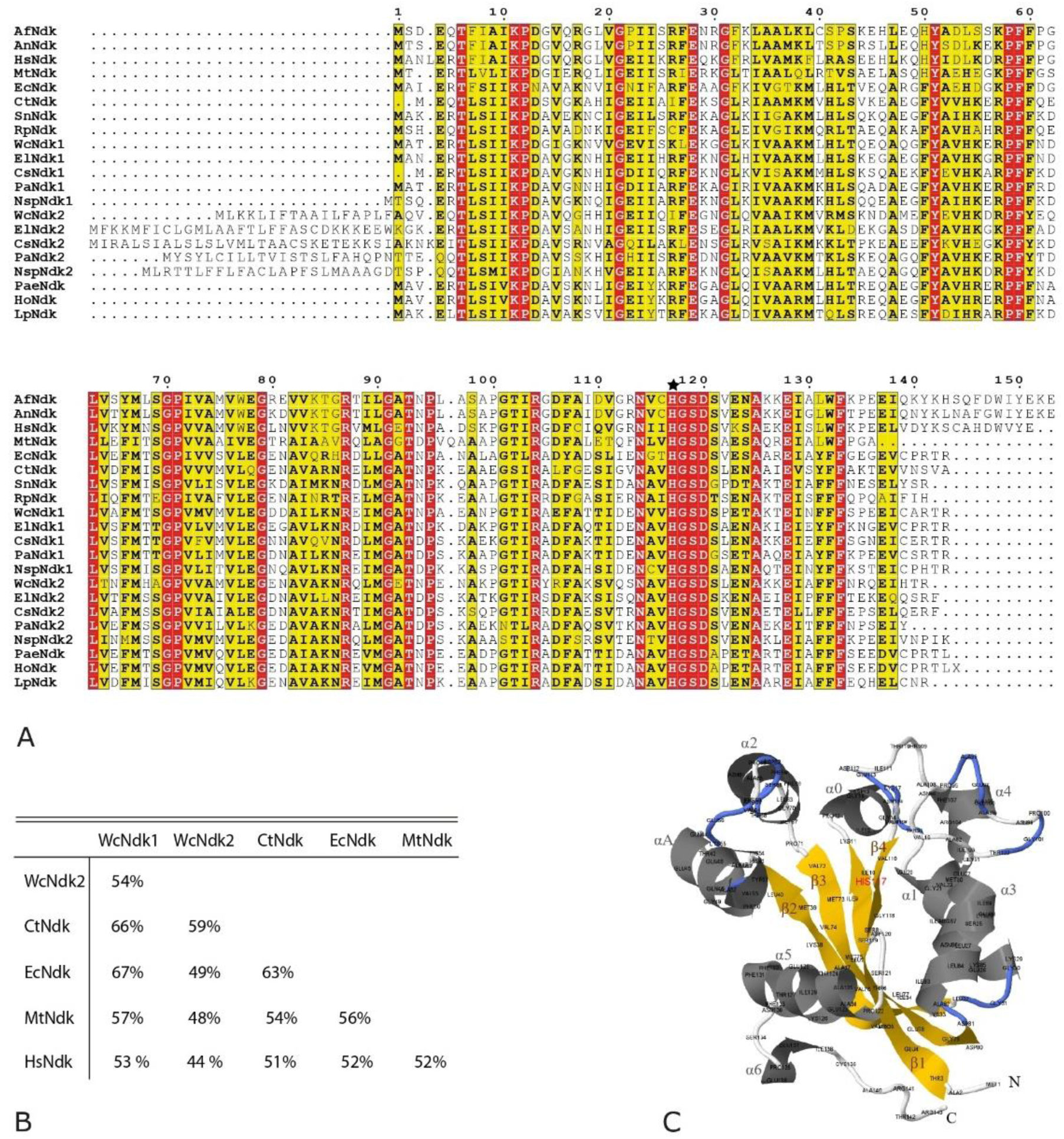
Ndk sequence and structure conservation. (A) Multiple sequence alignment of Ndk proteins across eukaryotes and prokaryotes. Conserved residues are highlighted in red with the highly conserved H residue, essential for catalytic activity, marked with an asterisk. Af: *Aspergillus flavus*, An: *A. niger*, Hs: *Homo sapiens*, Mt: *Mycobacterium tuberculosis*, Ec: *Escherichia coli,* Ct: *C. trachomatis*, Sn: *Simkania negevensis*, Rp: *Rhabdochlamydia porcellionis,* Wc: *W. chondrophila*, El: *Estrella lausannensis,* Cs: *Criblamydia sequanensis,* Pa: *Parachlamydia acanthamoebae,* Nsp: *Neochlamydia sp.*, Pae: *Pseudomonas aeruginosa*, Ho: *Halofilum ochraceum*, Lp: *Legionella pneumophila* (B) Amino acid sequence similarity matrix comparing Ndk proteins from *W. chondrophila* (WcNdk1 and WcNdk2), *Chlamydia trachomatis* (CtNdk), *Escherichia coli* (EcNdk), *Mycobacterium tuberculosis* (MtNdk), and human Ndk (HsNdk). (C) Predicted 3D structure of WcNdk1, illustrating its conserved α/β fold characteristic of the Ndk family. 3D structure reconstructed using Phyre2 and visualized using Jmol v.16.2.15. The H177, highlighted in red, is a critical histidine residue and positioned within the active site.

*W. chondrophila* contains two copies of *ndk*: *wcndk1 (*locus *wcw_1543*) and *wcndk2* (locus *wcw_1545*), with a sequence similarity of 54% (Figure 2B). The most striking difference between the two *W. chondrophila ndk* paralogs is the presence of a predicted signal peptide of 17 amino acids in WcNdk2, which is absent in WcNdk1.

The sequence similarity matrix of *W. chondrophila* Ndk proteins, WcNdk1 and WcNdk2, and their counterparts from other bacterial and eukaryotic species shows differing degrees of conservation as shown in Figure 2B. WcNdk1 shares the highest sequence identity with *E. coli* Ndk (EcNdk) and *Chlamydia trachomatis* Ndk (CtNdk) (67 % and 66%, respectively). In contrast, WcNdk2 exhibits slightly lower sequence similarity across species, which may indicate functional divergence of the two Ndks following gene duplication. Interestingly, both *W. chondrophila* Ndk homologs display relatively high sequence identity (∼50%) with human Ndk (HsNdk), which suggests the conservation of core functional domains across vast evolutionary distance. Sequence similarity matrix of Ndk proteins across different species is shown in Table S2.

WcNdk1 has 143 residues forming a polypeptide chain and adopting a very similar fold to Ndk protein from other origins including human Ndk^24,25^ (Figure 2C). One Ndk unit has α/β domains comprising a four-stranded antiparallel β-sheet and two connecting α-helices. This high degree of sequence and structural conservation between eukaryotes and the pathogens may enable them to interfere with host cellular process by mimicking host Ndk.

### Expression profile of *W. chondrophila* Ndks during infection

Temporal expression levels of genes and proteins across different developmental stages may help us understand their stage-specific roles and regulatory mechanisms. Since each of these stages has unique biological features, we can indirectly infer a protein’s function by mapping both transcript and protein levels across these developmental stages.

We quantified the expression profiles of *W. chondrophila ndks*, at both the transcript and protein level during McCoy cell infection (Figure 3). *wcndk1* and *wcndk2* transcripts were highly abundant at early time points (3 - 8 hpi) as revealed by RT-qPCR, then the mRNA level declined sharply to a minimum at 32 hpi, with a moderate surge toward the end of the cycle (48 hpi) (Figure 3A). In parallel, protein expression profile of WcNdk1 or WcNdk2 were analyzed using antibodies specifically raised against each protein. No WcNdk1 or WcNdk2 protein could be detected by Western blot of the *W. chondrophila*-infected cells at 3 to 8 hpi (Figure 3B, C). Both proteins became detectable by 24 hpi by Western blot and reached their highest signal intensity at this time point after normalization to bacterial number, followed by a drop to roughly 30% of that level by 48 hpi. To further assess protein expression at early stages, we performed immunofluorescence microscopy. In contrast to Western blot results, WcNdk1 and WcNdk2 signals were already detectable from 0 to 16 hpi, colocalizing with *W. chondrophila* (Figure S1). Together, these results suggests that both WcNdk1 and WcNdk2 are expressed throughout the developmental cycle, with an early transcriptional peak. While protein expression begins early, overall accumulation or detection sensitivity by Western blot peaks around 24 hpi, followed by a decline toward the end of the cycle.

**Figure 3:**
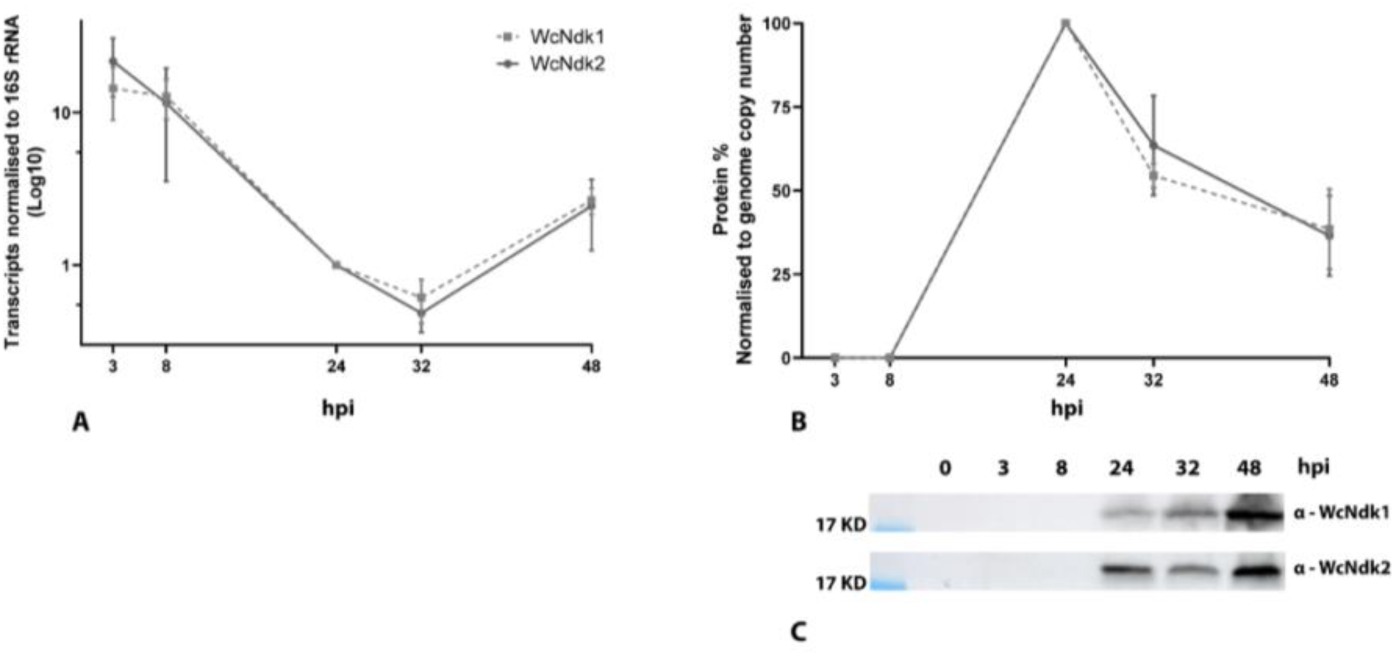
Temporal expression analysis of *ndk* paralogs in *W. chondrophila* during infection. (A) Quantitative RT-PCR analysis of *ndk1* and *ndk2* transcript levels at different hours post-infection (hpi), normalized to 16S rRNA. Transcript abundance is expressed relative to the 24 hpi time point, which was set as the reference (1.0). (B) Relative protein levels of WcNdk1 and WcNdk2 quantified by Western blot, normalized to bacterial genome copy number. Protein expression is shown relative to 24 hpi, set as 100%. (C) Representative Western blot images showing WcNdk1 and WcNdk2 expression across the infection time course. Data are shown as mean ± SD of 3 independent experiments.

### Subcellular localization of WcNdk1 and WcNdk2 in *C. trachomatis* heterologous expression

While signal peptides are typically associated with protein secretion, previous studies have demonstrated that Ndk can be secreted in certain bacterial species despite lacking a canonical signal sequence^26–28^. Moreover, the secretion of both WcNdk1 and WcNdk2 into the host cell cytoplasm is experimentally shown^29^, suggesting that WcNdk2 signal peptide may not contribute to secretion. The presence of a signal peptide on WcNdk2 suggests that it may be translocated to a specific subcellular compartment, giving it a biological role distinct from WcNdk1. To assess this possibility, we overexpressed a V5-tagged versions of WcNdk1 and WcNdk2 in *C. trachomatis* using the inducible shuttle vector pGL2 (Unpublished) and studied their subcellular localization by immunofluorescence and confocal microscopy (Figure 4). In this heterologous expression assay, WcNdk1 was exclusively detected inside the inclusion, closely associated with *C. trachomatis*, and no signal was observed in the host cell cytoplasm. Although this does not rule out secretion of WcNdk1, it suggests limited accumulation or detection outside the inclusion. In contrast, WcNdk2 was detected within both chlamydial inclusions and the host-cell nucleus, suggesting its trafficking into the nucleus. To further investigate the nuclear localization of WcNdk2, we cloned the gene into the pBOMBL vector^30^ and overexpressed the protein in *C. trachomatis*. Upon infection of McCoy cells with the transformed bacteria, inclusions were smaller, and bacterial growth was impaired compared to controls. Due to these growth defects, reliable localization of WcNdk2 could not be realized in this system. This vector-associated toxicity likely reflects bacterial stress caused by excessive or mis-regulated expression of WcNdk2, similar to the overexpression-associated toxicity reported for type III effectors in *Pseudomonas aeruginosa*^31^. Consistently, WcNdk2 expression was also not detected in the *Yersinia enterocolitica* type III secretion assay, suggesting poor tolerance or instability of the protein in this system.

**Figure 4:**
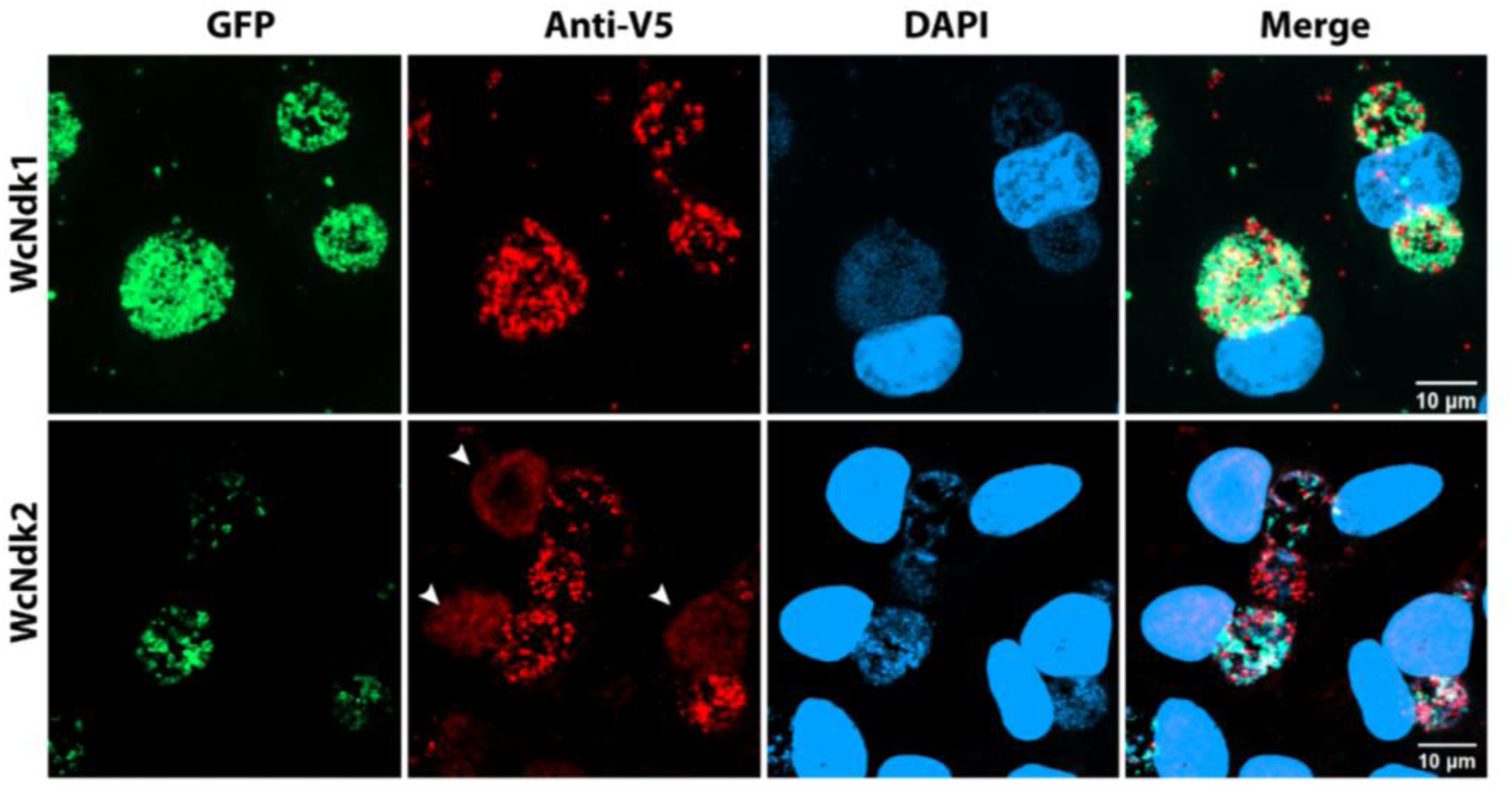
Subcellular localization of WcNdk1 and WcNdk2 overexpressed in *C. trachomatis. C. trachomatis* was transformed with plasmids expressing either WcNdk1 or WcNdk2 tagged with a V5 epitope. Infected McCoy cells were fixed at 24 hpi. GFP (green) indicates transformed *C. trachomatis*, anti-V5 (red) detects WcNdk proteins, and DAPI (blue) stains host cell nuclei and bacterial DNA. White arrows point to V5 signal within host nuclei.

### Subcellular localization of WcNdk1 and WcNdk2 in transfected HEK293T cells

To further confirm the observed subcellular localization of WcNdk1 and WcNdk2 in another heterologous system, we transiently expressed the V5-tagged WcNdk1 and WcNdk2 in HEK293T cells and analyzed their distribution by immunofluorescence microscopy. WcNdk1 was found exclusively in the cytoplasm and was absent from cell compartments such as nucleus or Golgi (Figure 5). WcNdk2 showed two mutually exclusive distribution patterns. In some cells the V5 signal was found throughout the nucleus, while in others, it was localized to perinuclear-Golgi regions, with no nuclear signal (Figure 5). A similar localization of WcNdk2 was also observed in transfected HeLa cells (data not shown), indicating that this targeting is not cell-type specific.

**Figure 5:**
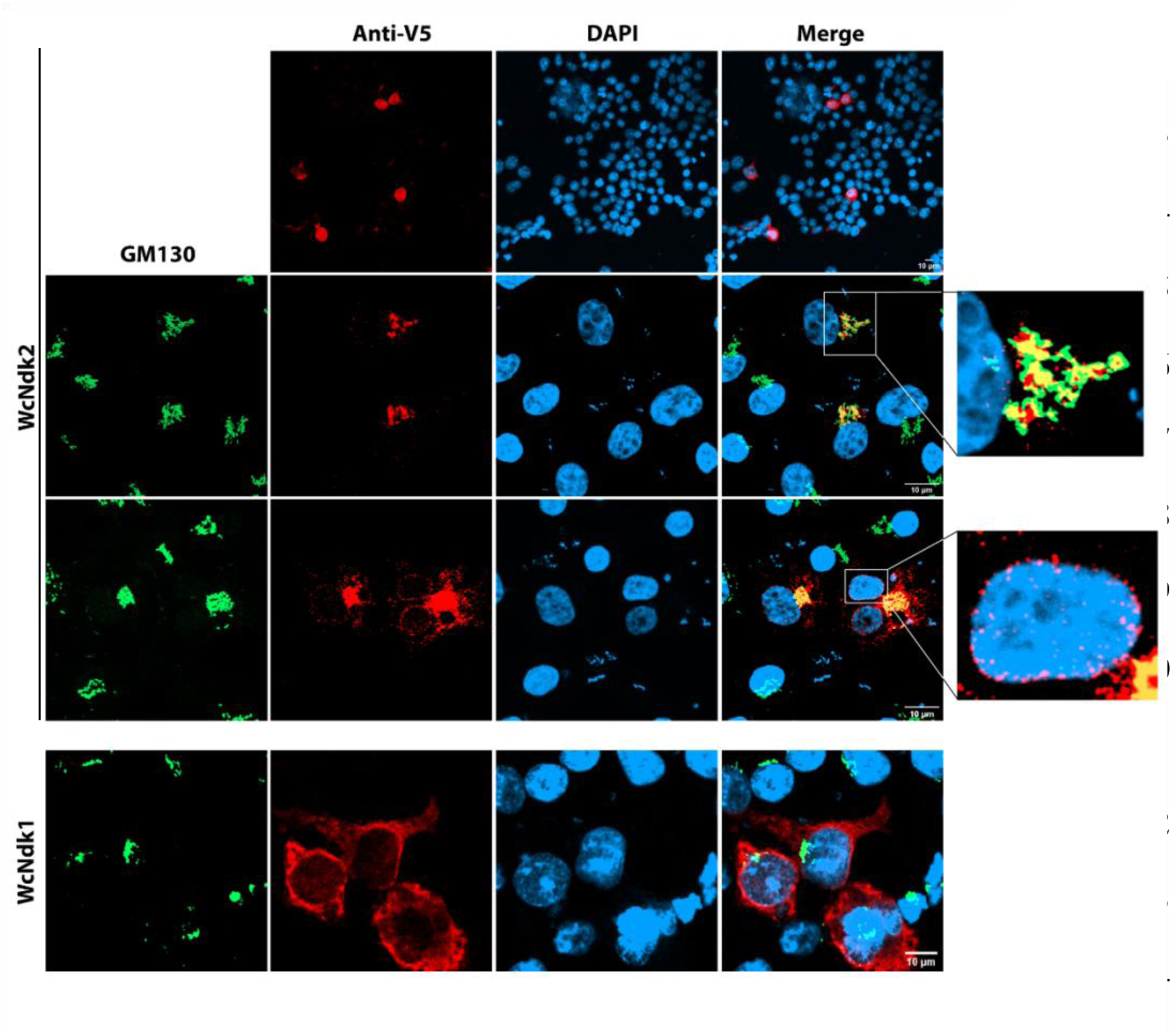
Subcellular distribution of WcNdk1 and WcNdk2 in HEK293T. HEK293T cells were transiently transfected with C-terminal V5-tagged WcNdk1 and WcNdk2 and stained with the Golgi marker GM130 (green), anti-V5 antibody (red), and DNA (DAPI, blue). High-magnification insets highlight the co-localization of WcNdk2 (red) with the Golgi (green) or the perinuclear region.

### The Ndk inhibitor, Azidothymidine (AZT), inhibits the growth of *W. chondrophila*

Since *W. chondrophila* is currently genetically intractable, we employed chemicals to inhibit Ndk function in this pathogen. AZT (3′-azido-3′-deoxythymidine), also known as zidovudine, is a known Ndk inhibitor^14,32–36^. This pharmaceutical, primarily approved for the treatment of human immunodeficiency virus (HIV), inhibits viral reverse transcription through chain termination^37^. This compound is structurally a thymidine analogue, where the 3’-hydroxyl group on the deoxyribose sugar is replaced by an azido group (N_3_) (Figure 6A).

**Figure 6:**
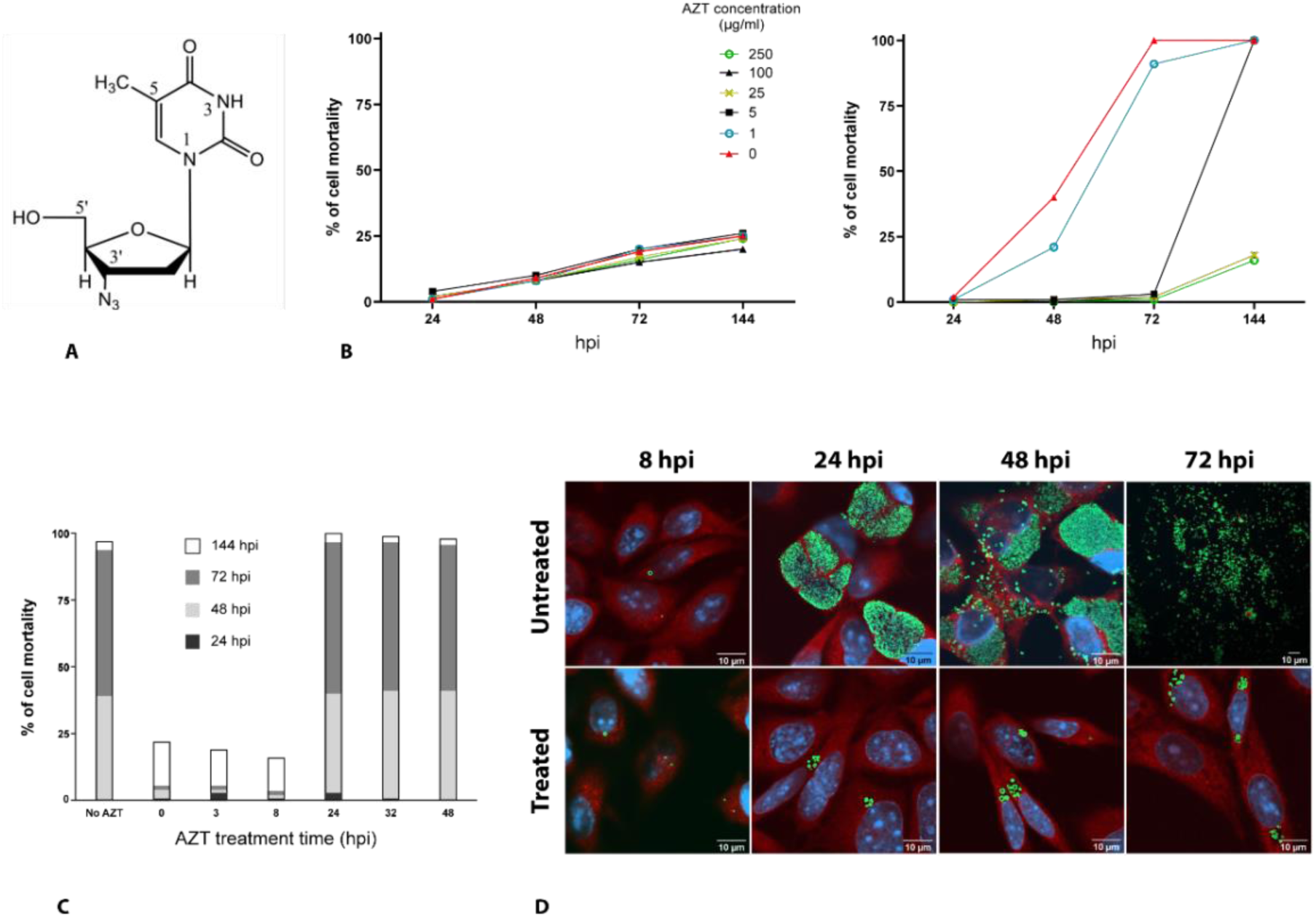
AZT inhibits *W. chondrophila* growth in infected host cells. (A) Chemical structure of azidothymidine (AZT). (B) AZT treatment of uninfected host cells, serving as a negative control (left panel) and treatment of infected host cells at concentrations ≥25 μg/mL (right panel). (C) Effect of AZT addition on *W. chondrophila*-infected cultures at various time points after infection. (D) Confocal microscopy of *W. chondrophila*-infected cells treated or not treated with AZT. In untreated conditions, *W. chondrophila* formed typical intracellular inclusions, whereas AZT treatment resulted in enlarged, aberrant structures. Bacteria are labeled using anti-*W. chondrophila* antibody (green); host cell nuclei are stained with DAPI (blue); and the host structures are marked with Concanavalin A (red).

The inhibition of the Ndk from *Aspergillus flavus* by AZT is documented, where AZT forms a strong hydrogen bond with key active site residues of Ndk (Arg-104, His-117 and Asp-120) and inhibits its enzymatic activity^14^. Since these three residues are highly conserved in Ndks of diverse organisms, including WcNdk1 and WcNdk2, we hypothesized that AZT could also serve as an effective inhibitor of Ndks in *W. chondrophila* (Figure 2A).

To test this hypothesis, we treated both uninfected and *W. chondrophila*-infected McCoy cells with increasing concentrations of AZT following infection and measured cell death resulting from bacterial proliferation using propidium iodide (PI) staining. In the absence of AZT or at low concentrations (1 µg/ml), *W. chondrophila*-infected cultures exhibited a sharp increase in cell mortality over time, reaching maximal levels by 144 hpi. Treatment with AZT at concentrations above 25 µg/ml significantly suppressed infection-induced cell death. A delay in cell mortality was observed with the intermediate concentration of 5 µg/ml (Figure 6B, right panel). In contrast, in uninfected cells, cell lysis remained comparable across all AZT concentrations, showing no significant difference from the no-AZT control. Therefore, the slight increase in cell death observed over time in untreated cultures is due to normal cell aging and turnover, and not due to AZT (Figure 6B, left panel). These results indicate that AZT did not exhibit significant adverse effects on mammalian host cells and provided a protective effect against *W. chondrophila*-induced cell death in a dose-dependent manner.

### Only early AZT treatment prevents *W. chondrophila*–mediated host cell death

To define the temporal window in which AZT exhibits its inhibitory effect on *W. chondrophila* growth, *W. chondrophila*-infected McCoy cell monolayers were treated with 25 µg/ml AZT at 0, 3, 8, 24, 32, or 48 hpi and the propidium iodide (PI) uptake was monitored at 24, 48, 72, and 144 hpi (Figure 6C). When AZT was added at 0, 3, or 8 hpi, cell mortality remained minimal (approximately 25% at 144 hpi), comparable to uninfected controls, indicating complete protection against bacterial-induced cell death. In contrast, administering AZT at 24 hpi or later failed to prevent cell death, resulting in maximal mortality, comparable to the no-AZT control. This indicates that once RB replication and inclusion expansion are established, AZT cannot reverse the course of infection.

### AZT induced the production of aberrant bodies (ABs) in *W. chondrophila*

To investigate the cellular-level effects of AZT treatment on *W. chondrophila* infection, we performed confocal microscopy on *W. chondrophila*-infected McCoy cells treated with 25 µg/ml AZT at 0 hpi. In untreated cultures, *W. chondrophila* underwent normal intracellular development, leading to host cell lysis at the end of infection. At 8 hpi, *W. chondrophila* EBs were observed attached to the host cell surface and internalized in both AZT-treated and untreated cultures indicating that AZT does not interfere with bacterial attachment or entry (Figure 6D). By 24 hpi, inclusions had formed in both conditions. However, in AZT-treated cells, these inclusions failed to expand, and only a limited number of bacteria were detected within them. This suggests that although EB-to-RB differentiation and initial rounds of replication may occur, bacterial proliferation is subsequently arrested. In addition, the *W. chondrophila* RBs appeared slightly enlarged in AZT-treated cultures, consistent with the formation of aberrant bodies (ABs). These abnormal conditions persisted until 72 hpi, indicating a disruption of the normal developmental cycle likely resulting in bacterial persistence.

### Effect of AZT on *W. chondrophila* growth is probably correlated with WcNdk2 activity

*W. chondrophila* encodes two *ndk* paralogs, organized in a single operon, with *pabA* positioned between them. To investigate which *ndk* copy mediates susceptibility to AZT, and whether this susceptibility is linked to the presence of the *pabA* gene, we selected representative *Chlamydiota* species with different *ndk* operon configurations and compared their response to AZT treatment. The selected species include *C. trachomatis*, which carries a single *ctndk* gene and lacks *pabA*; *Simkania negevensis,* which also has a single *snndk* gene but retains *pabA* elsewhere in the genome; *Estrella lausannensis*, which harbors a two-gene *elndk1*–*elndk2* operon without *pabA*; and *W. chondrophila*, which contains the full *wcndk1*–*pabA*–*wcndk2* operon (Figure 7, bottom panel). To evaluate the effect of AZT on these species, McCoy cell monolayers were infected, treated with 25 µg/ml at 0 hpi and fixed at 48 hpi before immunostaining and observation under confocal microscopy. As shown in Figure 7C, *trachomatis* and *S. negevensis* exhibited normal intracellular development in presence of AZT, indicating no significant growth inhibition. In contrast, *E. lausannensis*, like *W. chondrophila*, displayed enlarged intracellular structures resembling aberrant bodies. This suggest that AZT inhibitory effect depends on the presence of *ndk2*, regardless of the presence of *ndk1* and *pabA*.

**Figure 7:**
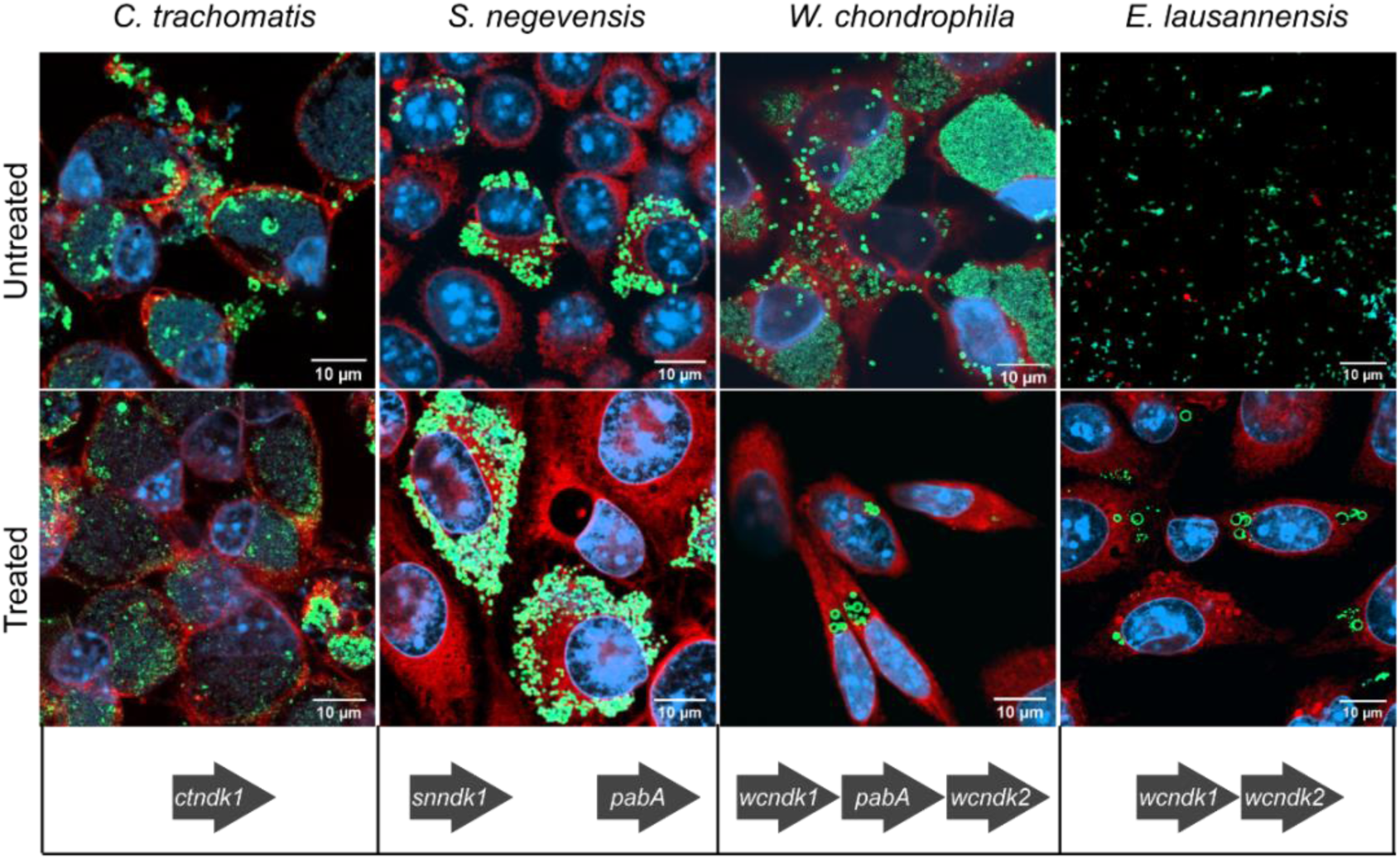
AZT sensitivity is restricted to *Chlamydiota* species encoding *ndk2*. Representative confocal micrographs of McCoy cells infected with *C. trachomatis*, *S. negevensis*, *W. chondrophila* or *E. lausannensis*, treated or not treated with 25 μg/ml AZT. The infected cells were fixed and stained 48 hpi. The lower panel shows the *ndk* operon arrangement: *C. trachomatis* and *S. negevensis*, each carry a single *ndk* gene. *S. negevensis* retains *pabA* elsewhere in the genome. *W. chondrophila* carries the two *wcndk* genes with a *pabA* in the middle. *E. lausannensis* encodes a two-gene *elndk1–elndk2* operon but lacks *pabA*. Bacteria are stained with species-specific antibodies. Host cytoplasm is labeled with Concanavalin A (red), and nuclei are stained with DAPI (blue). *ctndk*: *C. trachomatis ndk1*, *snndk1*: *S. negevensis ndk1*, *elndk1*: *E. lausannensis ndk1*, *elndk2*: *E. lausannensis ndk2*.

## DISCUSSION

The diversity of *ndk* copy number within members of the *Chlamydiota* reveals lineage-specific adaptations with potential metabolic and pathogenic consequences. While most *Chlamydiota* species possess a single *ndk* gene, a subset, including members of the *Parachlamydiaceae*, *Waddliaceae*, and *Criblamydiaceae*, harbor a second copy (*ndk2*), likely arising from a gene duplication event. This duplication may have provided a selective advantage by providing increasing metabolism flexibility, which could, in part, account for the comparatively faster growth of *W. chondrophila* and *E. lausannensis* in mammalian cells relative to other members of the *Chlamydiota* such as *Chlamydia* spp. and *S. negevensis*. Such an expansion of metabolic capacities might also represent an adaptation facilitating persistence and replication within free-living protists, suggesting that these lineages have evolved mechanisms enabling survival across diverse host environments. This is in line with their larger genome size (>2 Mb), expanded metabolic genes and their reduced dependence on host-derived metabolites. On the other side, the absence of the second copy of the *ndk* gene in *Rhabdochlamydiaceae*, *Simkaniaceae* and *Chlamydiaceae* coincides with their adaptations to more specific host niches and reduced metabolic pathways. The *pabA* gene is also absent in all members of the *Chlamydiaceae* family. In this family, enzymes from other biosynthetic pathways are recruited to meet their folate requirements^38^. The mechanisms driving the evolution of this operon in certain chlamydial families remains unknown. Further studies are needed to clarify the operon’s functional role and its evolutionary pathways in these families.

Both *W. chondrophila ndk* paralogs retain the universally conserved HGSD active-site motif and surrounding hydrophobic residues essential for autophosphorylation and phosphotransferase activities. This conservation suggests that both Ndk1 and Ndk2, despite their apparent different subcellular localizations, exert their functions, at least partially, through phosphorylation mechanisms.

The temporal expression patterns of *wcndk1* and *wcndk2* provide insights into their potential biological roles and regulation during the *W. chondrophila* developmental cycle. The organization of *wcndk1* and *wcndk2* within a single operon explains their synchronized expression profiles. The apparent mid-cycle surge in Ndk protein at 24 hpi coincides with the maximal bacterial replication, an increased demand for NTPs, and an increased host-manipulation activities that are essential for intracellular growth.

Despite sharing conserved kinase domains, our study showed significant differences in subcellular localization of WcNdk1 and WcNdk2. This divergence suggests functional specializations and may indicate that the two proteins utilize divergent cellular trafficking mechanisms during *W. chondrophila* infection.

In the *C. trachomatis* expression system, WcNdk1 was confined to the inclusion. This restricted distribution suggests that WcNdk1 may primarily serve bacterial intracellular functions within the inclusion. However, a previous study provided evidence for the secretion of WcNdk1 into the host cell cytosol^29^. It is, therefore, possible that WcNdk1 is secreted transiently or at levels insufficient to be detected via immunofluorescence. In contrast, WcNdk2 exhibited nuclear localization in *C. trachomatis* and HEK293T expression systems. This nuclear localization is consistent with previous studies on other pathogens, where Ndk was shown to bind host DNA and regulate gene expression through DNA cleavage^17,18^. Such putative nuclear activity suggests a potential role for WcNdk2 in reprogramming host transcriptional responses to favor bacterial survival or pathogenicity. It would be of great interest to identify specific host genes targeted by WcNdk2, as this could uncover previously unrecognized mechanisms by which *W. chondrophila* manipulates host cell function and could provide broader insights into its pathogenesis.

When expressed in HEK293T cells, WcNdk2 localized not only to the nucleus but also to the perinuclear-Golgi region. Bacterial Ndks, such as *Porphyromonas gingivalis* Ndk, are found in the perinuclear area of the host cells. This localization has been linked to the modulation of host purinergic signaling via P2X₇ receptors^28^, implying that the perinuclear localization of WcNdk2 could be a biologically relevant phenomenon rather than nonspecific aggregation. Unlike the perinuclear accumulation of bacterial Ndks, Golgi localization represents a novel observation not previously reported for any bacterial species. One plausible explanation for an association of WcNdk2 with the Golgi is that, after translocation into the host cytosol, WcNdk2 enters the classical ER–Golgi secretory pathway and is packaged into Golgi-derived vesicles for extracellular release to hydrolyze host extracellular ATP (eATP) and subvert purinergic signaling. Another possible explanation for the Golgi recruitment of WcNdk2 is that it might play a role in modulating vesicle trafficking between the Golgi and the bacterial inclusion. Although *W. chondrophila* does not redirect sphingomyelin transport from the Golgi to its inclusion^39^, it may still influence other aspects of vesicular trafficking, altering vesicle formation or fusion processes, to benefit bacterial survival or replication.

AZT is a well-established inhibitor of Ndk^14,32–36^. In our study, AZT treatment of *W. chondrophila*-infected cells protected them from bacteria-induced cell death and bacterial growth was arrested in the replication phase. Our comparative data across *Chlamydiota* species suggest that WcNdk2 is a potential target for AZT. WcNdk2 localizes to host cell compartments when expressed in heterologous systems and this positioning may increase its exposure to AZT. Thus, the growth arrest caused by AZT may result from disruption of Ndk2-dependent pathogenic mechanisms required for successful host-cell manipulation. Our results also showed that AZT must be applied before 24 hpi, to fully prevent *W. chondrophila*–induced cytotoxicity. This early phase of infection coincides with maximum RB replication, inclusion expansion, and active vesicular nutrient trafficking from the Golgi and ER^40^. In contrast, WcNdk1, which predominantly localizes within the bacteria, and is likely involved in maintaining bacterial nucleotide pools, appears less susceptible to AZT, possibly due to limited drug access to the inclusion compartment. These findings collectively support a model in which the inhibition of host-targeted, secreted WcNdk2, rather than the inclusion-confined WcNdk1, plays the key role in AZT’s antimicrobial activity against *W. chondrophila*.

Based on these observations, we propose a functional model for the two *W. chondrophila* Ndks, as illustrated in Figure 8. Together, these findings highlight the multifunctional nature of Ndk proteins and shed light on the distinct roles of *W. chondrophila* Ndks. However, the lack of genetic manipulation tools in *W. chondrophila* currently impedes direct investigation of Ndk function via gene deletion or mutagenesis. Although localization studies using heterologous expression systems suggest that WcNdk2 may interact with host cells, the possibility of overexpression artifacts or mis-localization cannot be entirely ruled out. Additionally, although AZT is a well-established Ndk inhibitor with no other known bacterial targets, direct in vitro evidence confirming WcNdk2 as the AZT target is still missing. In vitro ATPase assays are essential to validate this interaction. In the absence of such studies the proposed functions remain speculative. The advancement of genetic tools would be essential to validate the proposed roles of WcNdk1 and WcNdk2 in *W. chondrophila* metabolism and host interaction.

**Figure 8:**
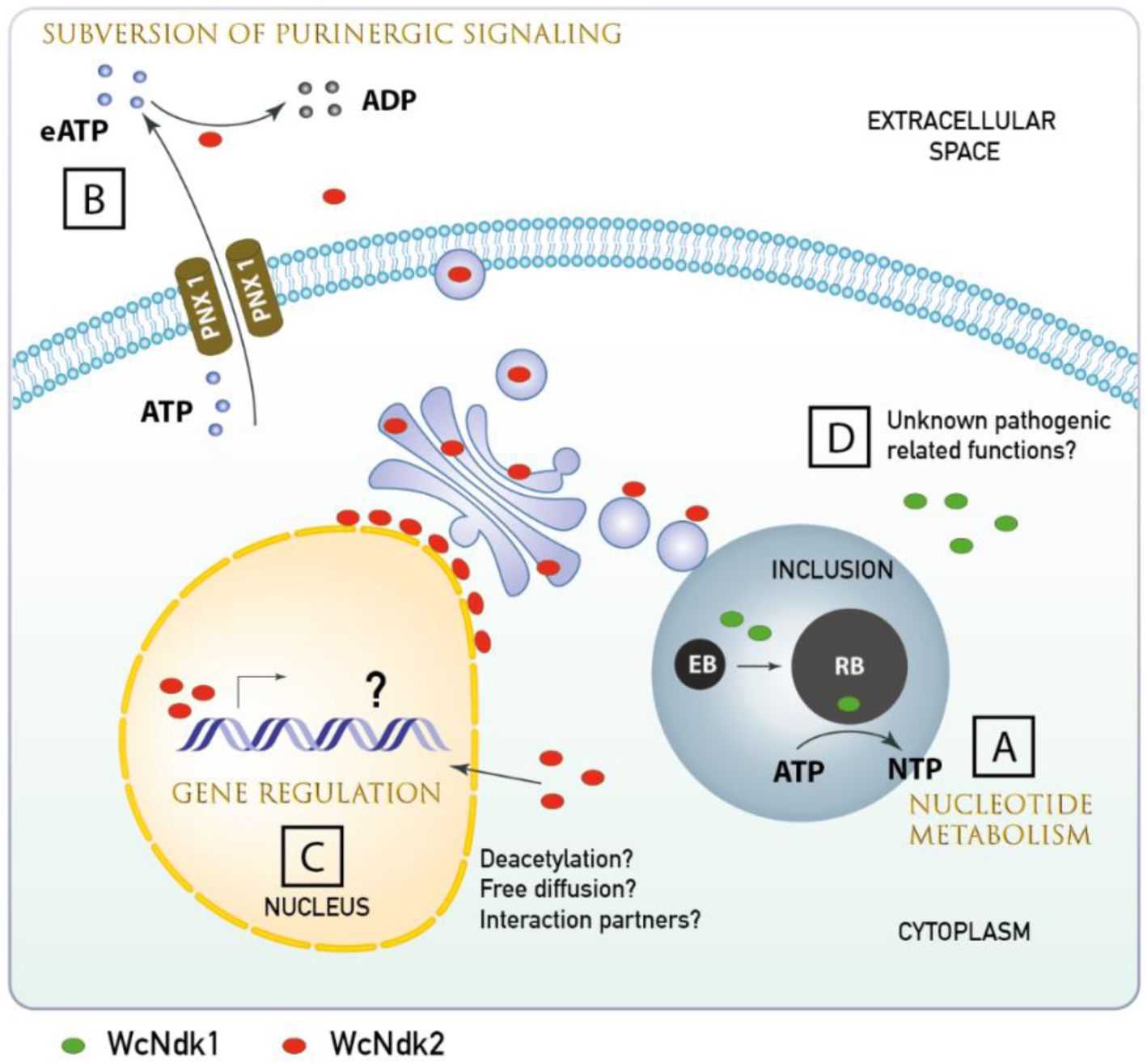
Proposed model for the functions of WcNdk1 and WcNdk2 in *W. chondrophila.* This schematic illustrates the distinct subcellular localizations and putative roles of WcNdk1 (green) and WcNdk2 (red), during infection of a host cell. A: WcNdk1 is predominantly retained within the bacterial inclusion, where it sustains basic nucleotide metabolism and “housekeeping” functions. B: A fraction of WcNdk1 is secreted across the inclusion membrane into the host cytosol, however, the specific functions of cytosolic WcNdk1 remain undefined. C: WcNdk2 is observed accumulating within the host nucleus. Its import into the nucleus could occur via one or more of the following mechanisms: a deacetylation event allowing passive nuclear import; free diffusion through nuclear pores owing to its small molecular size; or interaction with host proteins that escort WcNdk2 into the nucleus. Inside host nucleus, WcNdk2 could bind directly to DNA leading to up- or downregulation of host genes involved in immunity or apoptosis. The precise gene targets remain unknown. D: WcNdk2 traffics through Golgi/ER-derived vesicles and perinuclear region. This localization may be related to the packaging of WcNdk2 into vesicles destined for secretion to subvert host purinergic signaling through hydrolysis of eATP into ADP. E: The association of WcNdk2 with the Golgi apparatus may imply an involvement in vesicle trafficking and acquisition of host-derived metabolites necessary for expansion of the inclusion and bacterial survival. Illustration generated using Adobe Illustrator 2024 (Adobe Inc., USA).

## CONCLUSION

Given its diverse activities, Ndk has attracted interest across multiple disciplines, including microbiology, cell biology, and drug development. In this study, we introduced chlamydial Ndk as a multifunctional protein with roles extending beyond nucleotide metabolism, highlighting its potential involvement in host–pathogen interactions. These findings not only enhance our understanding of Ndk biology in the *Chlamydiota* phylum but also point to Ndk2 as a potential therapeutic target, opening new avenues for dissecting host manipulation strategies in obligate intracellular bacteria.

## MATERIALS AND METHODS

### Phylogenetic and structural analysis

Species-based and *ndk* gene phylogenies were retrieved from the *Chlamydia* Database^41^. Trees were trimmed using FigTree (v1.4.4) to retain only relevant *Chlamydiota* species. The Ndk protein sequences across different taxa were retrieved from the National Center for Biotechnology Information (NCBI). Multiple sequence alignment of Ndk proteins across eukaryotes and prokaryotes was performed using the ClustalW tool in UGENE v1.30.0. The corresponding similarity heat map (percentage identity, excluding gap) was also generated on UGENE. Alignment visualization was carried out with EPSript https://espript.ibcp.fr/ESPript/cgi-bin/ESPript.cgi. The 3D structure of the protein was reconstructed using Phyre2^42^ and visualized using Jmol v.16.2.15.

### Cell culture and *W. chondrophila* infection

McCoy (murine fibroblast cells; ATCC CRL-1696, purchased from ATCC), HEK293T (ATCC CRL-11268, USA, obtained as a gift from Dr. Thierry Roger, Lausanne University Hospital) or HeLa cells (human cervical adenocarcinoma epithelial cells; ATCC CCL-2, a gift from Dr. Thierry Roger) were maintained at 37°C in 5% CO_2_ in DMEM GlutaMAX (Thermo Fisher Scientific, USA), supplemented with 10% fetal bovine serum (FBS; Thermo Fisher Scientific, USA). *W. chondrophila* (ATCC VR-1471) was propagated in *Acanthamoeba castellanii* (ATCC 30010) at 25°C in T-25 flasks containing 6 ml of peptone–yeast–extract–glucose broth. At the time of infection, lysed *W. chondrophila*-infected amoebae were filtered through a 5-µm syringe filter to remove amoebal debris. The bacterial solution was then used to infect the host cells at a dilution which was optimum for infection (MOI 0.1–1). To synchronize the infection, the infected cells were centrifuged at 1790 g for 10 minutes. They were then incubated at 37°C in 5% CO_2_ for 15 minutes. Following incubation, the inoculum was then replaced with fresh medium.

### Purification of recombinant His-tagged WcNdk1 and WcNdk2 for antiserum production

The *E. coli* strain BL21 containing pET28a-*wcndk1-6xHis* or pET28a-*wcndk2-6xHis* was grown to an OD_600_ of 0.5. The culture was induced using 1 mM Isopropyl β-D-thiogalactopyranoside (IPTG, Applichem, Germany) and incubated at 37°C for 4 hours to express the recombinant protein. Bacterial pellets were resuspended in native lysis buffer (50 mM NaH_2_PO_4_, 300 mM NaCl, and 10 mM imidazole, pH 8). The resuspended cells were lysed using a combination of methods: three cycles of freeze (ethanol-dry ice bath) and thaw (at 37°C), followed by chemical lysis with 1 mg/ml lysozyme (AppliChem, Germany) and sonication. The recombinant protein was purified under native condition using Ni-NTA agarose beads (Qiagen, Germany) according to the manufacturer’s instructions. Briefly, bacterial lysates were incubated with Ni-NTA resin for 2 hours at 4 °C with gentle rotation. The protein-bound resin was then loaded onto poly-prep chromatography columns (Bio-Rad, USA), washed with wash buffer (50 mM Tris-HCl, 300 mM NaCl, 20 mM imidazole, pH 8.0), and eluted with elution buffer containing 250 mM imidazole. The purified protein was dialyzed using Slide-A-Lyzer Dialysis Cassette, 2,000 MWCO (Thermo Scientific, USA) overnight against PBS to remove imidazole. The concentration of the protein was determined using Bradford’s reagent with BSA (Bio-Rad, USA) as a standard. Following protein purification, rabbit polyclonal antisera were produced using immunization services offered by Eurogentec SA (Seraing, Belgium).

### Western Blot

McCoy cells were seeded at a density of 1 × 10^6^ per T-25 flask one day before infection and infected with *W. chondrophila* as described above. Infected cells were harvested by scraping at the specified hpi and a fraction of the culture was saved for extraction and quantification of the genomic DNA (see below). The cell suspension was pelleted at 1790 g for 10 minutes, then washed twice with PBS, and finally resuspended in 0.5 ml of 1x Laemmli sample buffer (Bio-Rad, USA). An equal volume of each lysate was resolved on 12% SDS-PAGE precast gels (Bio-Rad, USA) and transferred onto an Amersham Protran nitrocellulose membrane (Cytiva, USA). The membrane was blocked in saturation buffer (10 mM Tris-Base, 150 mM NaCl, and 0.05% Tween 20) containing 5% nonfat dried milk (AppliChem, Germany) for 2 hours at room temperature (RT). Blots were probed overnight at 4°C with antibodies against WcNdk1 and WcNdk2 diluted in saturation buffer with 0.5% non-fat dried milk. After three washes with saturation buffer containing 0.5% non-fat dried milk, blots were incubated with secondary antibodies (horseradish peroxidase-conjugated anti-rabbit IgG; Promega, USA, or anti-mouse IgG; Bio-Rad, USA) for 1 hour at Room temperature and processed using the Amersham ECL detection system (Cytiva, USA). Western blot bands intensities were quantified using EvolutionCapt edge software (Vilber, France) and normalized to the corresponding bacterial genome copy number. Graphs were generated using GraphPad v. 10.4.1.

### Extraction of genomic DNA and quantification of *W. chondrophila* genome copy number

Genomic DNA from collected samples was extracted using the Wizard SV Genomic DNA purification system (Promega, USA) according to the manufacturer’s instructions. The extracted DNA served as a template for qPCR with iTaq Universal Probes Supermix (Bio-Rad, USA) to quantify the *W. chondrophila* bacterial population. The 16S copy numbers were determined using a standard curve, which was generated from serial dilutions of a plasmid containing one copy of the 16S rRNA gene. The primer sequences^8^ are listed in Table S1.

### RNA extraction and cDNA synthesis

*W. chondrophila-*infected McCoy cell monolayers or uninfected control cells were harvested at indicated hpi by scraping and centrifugation (5000 g, 10 minutes). Following centrifugation cells were lysed directly in TRIzol Reagent (Invitrogen, USA). Total RNA was extracted by chloroform separation and recovery of the aqueous phase (12,000 g, 15 min, 4 °C). RNA was precipitated with isopropanol, washed with 75 % ethanol, air-dried, and resuspended in RNase-free water. To remove genomic DNA, samples were treated with RNase-free DNase I (Invitrogen, USA) following the manufacturer’s instructions. cDNA was synthesized using the GoScript Reverse Transcription System (Promega, USA) with random primers. Reverse transcription was performed at 42°C for 60 minutes, followed by enzyme inactivation at 70°C for 15 minutes. All cDNAs were diluted 1:5 prior to qPCR.

### RT-qPCR

The qPCR reactions were set up in a final volume of 20 μl containing 4 μl of cDNA template, the appropriate concentration of each primer, and 1X iTaq Universal SYBR Green Supermix (Bio-Rad, USA). All samples were run in duplicates, and no-template controls were included for each primer pair to assess non-specific amplification. The fluorescent reporter signal was normalized against the internal reference dye (ROX) signal. qPCR was carried out on QuantStudio 3 Real-Time PCR System (Applied Biosystems, USA) using the following thermal program. A single cycle of DNA polymerase activation for 3 min at 95 °C, followed by 45 amplification cycles of 15 s at 95 °C (denaturing step) and 1 min at 60 °C (annealing and extension step). Gene expression data were normalized to *W. chondrophila* 16 rRNA gene, which served as the internal reference gene. Quantitative data from qRT-PCR experiments were collected from at least three independent replicates. The primers used for RT-qPCR are listed in Table S1.

### Application of AZT on *W. chondrophila* culture and cell death assessment

McCoy cells were seeded at a density of 1 × 10^4^ cells per well in 96-well plates (Corning, USA). Cells were infected with *W. chondrophila* or left uninfected as a control. To monitor cell death, Propidium Iodide (PI, 7 µg/mL, Sigma-Aldrich, Germany) was added to the growth medium. AZT was purchased from TOCRIS (Cat. No.:4150) and was added to the wells at concentrations ranging from 0 to 250 µg/mL in six technical replicates per condition. Depending on the experimental design, AZT was added at the time of infection (0 hpi) or at later time points (3, 8, 24, 32, and 48 hpi). PI fluorescence was measured at specified time points post-infection using a FLUOstar Omega plate reader (BMG LABTECH, Germany; excitation: 540 nm; emission: 640 nm). To define 100% cell death, 0.1% Triton X-100 was added to control wells prior to PI reading.

### Antibodies, immunofluorescence assay, and confocal microscopy

Mouse anti-V5 antibody were purchased from Thermo Fisher Scientific (USA). Goat anti-*Chlamydia trachomatis* major outer membrane protein (MOMP) antibodies were obtained from Lifespan Bioscience (LS-C55983, USA). Polyclonal antibodies against *W. chondrophila*, *E. lausannensis*, and *S. negevensis* were homemade. Antibodies against GM130 (a cis-Golgi marker) were obtained from BD Biosciences (USA). Secondary anti-mouse, and anti-rabbit or anti-goat antibodies conjugated to Alexa Fluor 488 or 594, as well as Texas Red–conjugated concanavalin A, were purchased from AppliChem (Germany).

McCoy cells were cultured on glass coverslips placed in 24-well-plates and infected with various bacterial species. At the indicated time points, cells were fixed with ice-cold methanol or 4% paraformaldehyde (PFA) for 5 minutes at 4°C, followed by three washes with phosphate-buffered saline (PBS). Fixed cells were permeabilized and blocked for 30 minutes using a blocking solution containing 0.1% saponin, 10% fetal bovine serum (FBS), and 0.04% sodium azide in PBS. Blocked cells were incubated with primary antibodies for 2 hours at RT (anti-V5 diluted 1:5000, anti-MOMP diluted 1:500 and anti GM130 diluted 1:300). After three washing steps with blocking solution, cells were incubated for 1 hour with Alexa Fluor 488- or 598-conjugated secondary antibodies (1:1000 dilution) to label bacteria. DAPI (4′,6-diamidino-2-phenylindole; Thermo Fisher Scientific, USA) (1:3000 dilution) and Concanavalin A (1:50 dilution) were used to stain nuclei and carbohydrates, respectively. Cells were then washed three times with PBS and briefly rinsed with purified water before mounting. Coverslips were mounted on glass slides with Mowiol 4-88 (Sigma-Aldrich, USA) and stored in the dark until imaging using Zeiss (LSM 900, Germany) confocal laser scanning microscope.

### pGL2 and pBOMBL plasmid construction

The genes *wcndk1* and *wcndk2* were amplified from *W. chondrophila* genomic DNA using primers with 5′ overhangs compatible with the pGL2 (a kind gift from the Scott Hefty laboratory, University of Kansas) or pBOMBL (generously provided by Scot Ouellette laboratory, University of Nebraska Medical Center) backbone. All PCR products included a C-terminal V5 epitope tag to facilitate detection. PCR products were purified using the QIAquick PCR Purification Kit (Qiagen, Germany). The pBOMBL vector was linearized with EagI and KpnI (NEB, USA). The native *C. trachomatis*–derived pGL2 vector (11,786 bp; β-lactamase selection marker) was linearized by Age I digestion (NEB, USA) and recovered from agarose gels with the QIAquick Gel Extraction Kit (Qiagen, Germany). Purified inserts and linearized vectors were assembled via In-fusion cloning using 5X In-fusion Snap assembly Master Mix (Takara Bio, Japan). Assemblies were incubated at 50 °C for 15 min. Transformations were performed using *E. coli* dam⁻ dcm⁻ cells. Positive clones were screened by colony PCR, and successful insertions were confirmed by Sanger sequencing using both vector- and insert-specific primers.

### *C. trachomatis* transformation

Transformation of *C. trachomatis* was performed with minor modifications to previously described methods^43^.1 × 10^6^ McCoy cells were seeded in 6-well plates and cultured overnight. 2.5 × 10^6^ plasmid-free *C. trachomatis* serovar L2 (EBs), were resuspended in 300µl Tris-CaCl_2_ buffer (10 mM Tris, 50 mM CaCl2, pH 7.4), and incubated with 2 µg of sequence-verified plasmid DNA for 30 minutes at room temperature. 1 mL Hank’s balanced salt solution (HBSS; Gibco, Thermo Fisher Scientific, USA) was then added to each reaction. This mixture was added to McCoy cells in a 6-well plate after removing the medium. The infection was carried out by centrifugation at 400 × g for 15 minutes at room temperature and incubation at 37°C for 15 minutes. Then, the inoculum was removed, and cells were incubated with 2 mL of DMEM medium supplemented with 10% FBS for 8 hours at 37°C and 5% CO₂. After this incubation, the medium was replaced with DMEM containing 10% FBS, 1 µg/mL cycloheximide (Sigma-Aldrich, Germany), and penicillin G (0.6 mg/mL; Sigma-Aldrich, Germany) or spectinomycin (50 µg/mL; Sigma-Aldrich, Germany) to select for transformed bacteria. Cells were passaged every 48 hours, and the development of fluorescent inclusions were monitored until they were clearly observed and established. The transformed *C. trachomatis* was harvested and titrated by IFU assay and stored at −80°C. For experiments requiring induction, 50 ng/ml anhydrotetracycline (aTc, Sigma-Aldrich, Germany) was added at the time of infection, whereas, in control conditions, the inducer was excluded.

### Gateway cloning for mammalian expression in pDEST47

For generation of pDEST47 expression plasmids, attB-flanked primers were used to amplify the C-terminal v5-tagged *wcndk1* and *wcndk2*. Entry clones were generated by BP recombination using the Gateway BP Clonase II Enzyme Mix (Thermo Fisher Scientific, USA). The resulting pDONR201-*ndk-v5* entry clones were selected on Kanamycin (AppliChem, Germany) plates. Destination vectors were generated via LR recombination of the entry clones with pDEST47 using the Gateway LR Clonase II Enzyme Mix (Thermo Fisher Scientific, USA) following the supplier’s instructions. Final constructs were transformed and validated as described above, with plasmid DNA prepared from overnight cultures using the GeneJET Plasmid Miniprep Kit (Thermo Fisher Scientific, USA).

### Transient transfection of HEK293T and HeLa cells

Glass coverslips placed in 24-well plates were coated with poly-L-lysine (100 µg/mL; Sigma-Aldrich, Germany) for 30 min at room temperature, washed three times with PBS, and air-dried for 3 h. HEK293T cells were seeded on coated coverslips at 2 × 10⁵ cells per well and incubated overnight at 37 °C in 5 % CO₂. HeLa cells were seeded on uncoated coverslips at 2.5 × 10⁵ cells per well and incubated under the same conditions. Twenty-four hours post-seeding, cells were transfected with pDEST47 expression plasmids using the Lipofectamine 3000 Transfection Kit (Thermo Fisher Scientific, USA). according to the manufacturer’s instructions. Briefly, for each well, 1 µg plasmid DNA was diluted in 25 µL serum-free DMEM containing 1 µL P3000 reagent. Separately, 0.75 µL Lipofectamine 3000 reagent was diluted in 25 µL serum-free DMEM. The two mixtures were combined, incubated for 15 min at room temperature, and 50 µL of the transfection complex was added dropwise per well. Cells were fixed 24 hours post-transfection with PFA 4% for downstream immunofluorescence analysis as described above.

## AUTHOR CONTRIBUTIONS

G.G.S.: Conceptualization, Methodology, Investigation, Formal analysis, Visualization, Writing – original draft.

K.K.B.: Investigation, Validation, Resources, Writing – review & editing.

S.E.A.: Validation (pBOMBL-based experiment), Writing – review & editing.

G.G.: Supervision, Conceptualization, Project administration, Funding acquisition, Writing – review & editing.

