## Supplementary Material for "Functional insights into nucleoside diphosphate kinases encoded by two *ndk* paralogs in *Waddlia chondrophila*"

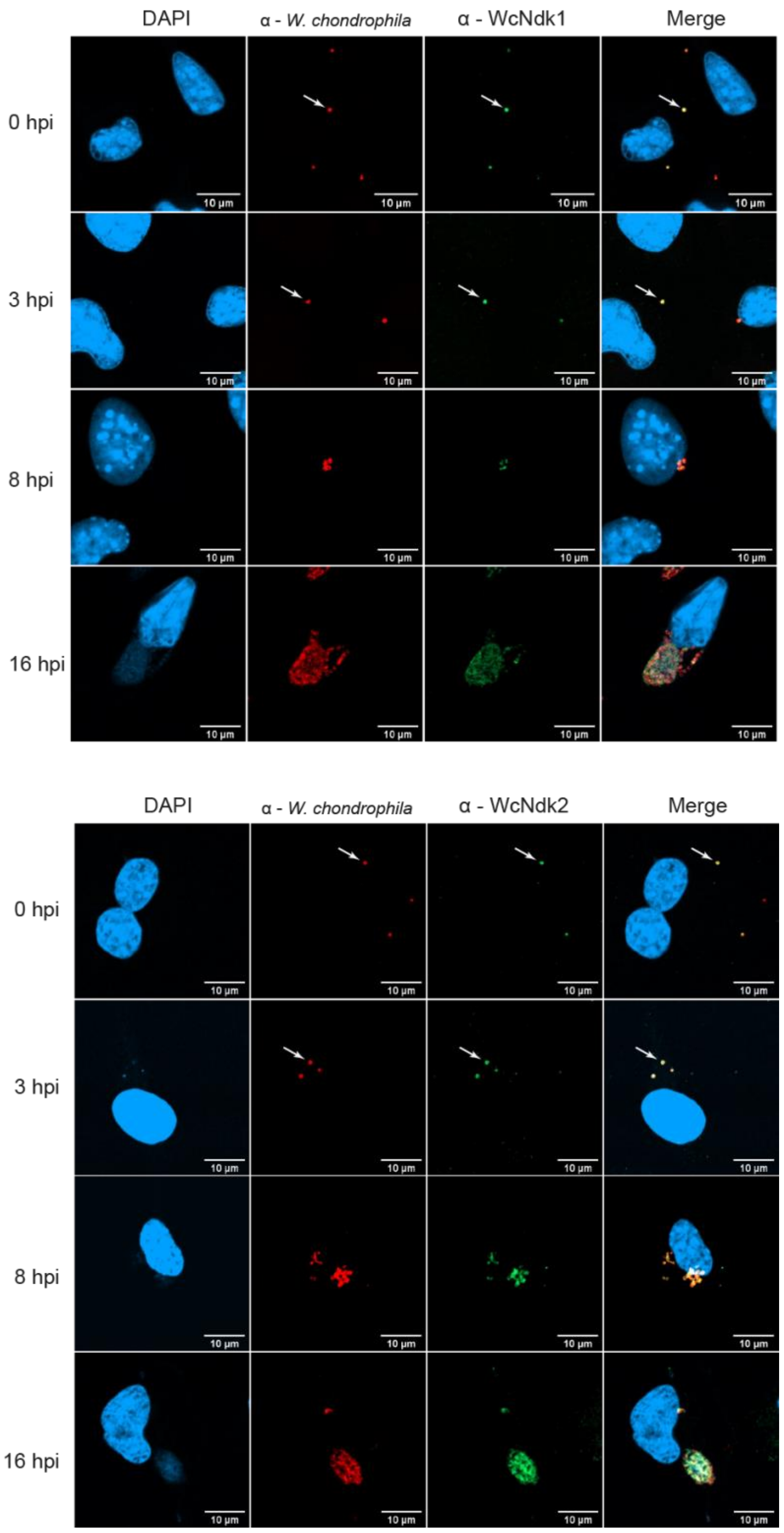


Figure S1: Immunofluorescence detection of WcNdk1 and WcNdk2 during *W. chondrophila* infection. DAPI staining (blue) marks host nuclei, anti-*W. chondrophila* (red) labels bacteria, and anti-WcNdk1 or anti-WcNdk2 (green) detect the corresponding proteins. Merged images show colocalization in yellow. Arrows indicate the localization of bacteria, WcNdks, and their colocalization.

| Target gene | Oligo type | Sequence (5' -> 3') | Label | Amplicon size (bp) |
|---|---|---|---|---|
| wcw_1543 (*wcndk1*) | Forward primer | tgg-tat-tgg-aag-gag-acg-atg-c | - | 160 |
| wcw_1543 (*wcndk1*) | Reverse primer | tcc-gtt-ttt-gcc-gtt-tct-gg | - | 160 |
| wcw_1545 (*wcndk2*) | Forward primer | agt-tgt-tgc-gat-ggt-act-gga | - | 150 |
| wcw_1545 (*wcndk2*) | Reverse primer | gaa-tct-gat-cca-tgc-acg-gc | - | 150 |
| *Waddlia* 16S rRNA | Forward primer | ggc-cct-tgg-gtc-gta-aag-ttc-t | - | 101 |
| *Waddlia* 16S rRNA | Reverse primer | cgg-agt-tag-ccg-gtg-ctt-ct | - | 101 |
| *Waddlia* 16S rRNA | TaqMan prob | FAM-cAt-ggg-aaC-aag-aga-agG-ATg-BHQ | FAM/BHQ-1 | - |

Table S1: Primer sequences used for RT-qPCR in this study. Sequence letters in capital are Locked Nucleic acids.

| | C. sequanensis_Ndk2 | E. lausannensis_Ndk2 | W. chondrophila_Ndk2 | Neochlamydia sp_Ndk2 | P. canthamoebae_Ndk2 | Neochlamydia sp_Ndk1 | C. sequanensis_Ndk1 | E. lausannensis_Ndk1 | P. acanthamoebae_Ndk1 | W. chondrophila_Ndk1 | Rabdochlamydia_Ndk | S. negevensis_Ndk | C. trachomatis_Ndk | P.aeruginosa_Ndk | Thioalbus dentrifivans_Ndk | Legionella pneumophila_Ndk | Halofilum ochreceum_Ndk | M. tuberculosis_Ndk | E. coli_Ndk | A.flavu_Ndk | A.niger_Ndk |
|---|---|---|---|---|---|---|---|---|---|---|---|---|---|---|---|---|---|---|---|---|---|
| C. sequanensis_Ndk2 | 100% | | | | | | | | | | | | | | | | | | | | |
| E. lausannensis_Ndk2 | 62% | 100% | | | | | | | | | | | | | | | | | | | |
| W. chondrophila_Ndk2 | 49% | 56% | 100% | | | | | | | | | | | | | | | | | | |
| Neochlamydia sp_Ndk2 | 48% | 52% | 57% | 100% | | | | | | | | | | | | | | | | | |
| P. canthamoebae_Ndk2 | 55% | 60% | 62% | 64% | 100% | | | | | | | | | | | | | | | | |
| Neochlamydia sp_Ndk1 | 51% | 56% | 55% | 57% | 57% | 100% | | | | | | | | | | | | | | | |
| C. sequanensis_Ndk1 | 50% | 56% | 62% | 56% | 60% | 77% | 100% | | | | | | | | | | | | | | |
| E. lausannensis_Ndk1 | 48% | 57% | 60% | 56% | 57% | 82% | 81% | 100% | | | | | | | | | | | | | |
| P. acanthamoebae_Ndk1 | 51% | 57% | 56% | 56% | 56% | 81% | 78% | 83% | 100% | | | | | | | | | | | | |
| W. chondrophila_Ndk1 | 49% | 52% | 54% | 59% | 53% | 76% | 73% | 79% | 79% | 100% | | | | | | | | | | | |
| Rabdochlamydia_Ndk | 46% | 50% | 57% | 53% | 52% | 65% | 64% | 66% | 66% | 66% | 100% | | | | | | | | | | |
| S. negevensis_Ndk | 51% | 55% | 57% | 54% | 57% | 77% | 73% | 77% | 78% | 76% | 69% | 100% | | | | | | | | | |
| C. trachomatis_Ndk | 47% | 53% | 59% | 53% | 55% | 71% | 70% | 72% | 70% | 66% | 67% | 69% | 100% | | | | | | | | |
| P.aeruginosa_Ndk | 47% | 52% | 56% | 54% | 54% | 72% | 70% | 78% | 76% | 80% | 68% | 77% | 66% | 100% | | | | | | | |
| Thioalbus dentrifivans_Ndk | 48% | 52% | 55% | 56% | 55% | 75% | 73% | 80% | 79% | 82% | 70% | 81% | 68% | 88% | 100% | | | | | | |
| Legionella pneumophila_Ndk | 52% | 54% | 59% | 55% | 58% | 75% | 72% | 76% | 74% | 72% | 69% | 79% | 72% | 77% | 80% | 100% | | | | | |
| Halofilum ochreceum_Ndk | 46% | 51% | 55% | 54% | 53% | 71% | 70% | 77% | 76% | 79% | 67% | 77% | 66% | 99% | 88% | 77% | 100% | | | | |
| M. tuberculosis_Ndk | 38% | 45% | 48% | 45% | 47% | 57% | 55% | 58% | 57% | 57% | 56% | 59% | 54% | 58% | 59% | 58% | 57% | 100% | | | |
| E. coli_Ndk | 43% | 48% | 49% | 48% | 49% | 66% | 66% | 72% | 68% | 67% | 65% | 68% | 63% | 70% | 71% | 68% | 69% | 56% | 100% | | |
| A.flavu_Ndk | 36% | 37% | 44% | 40% | 41% | 53% | 52% | 54% | 53% | 53% | 51% | 51% | 49% | 49% | 51% | 52% | 49% | 54% | 52% | 100% | |
| A.niger_Ndk | 36% | 37% | 43% | 39% | 41% | 51% | 51% | 52% | 52% | 53% | 49% | 52% | 51% | 49% | 51% | 52% | 49% | 54% | 51% | 89% | 100% |
| Homo sapiens_Ndk | 34% | 36% | 44% | 39% | 39% | 51% | 49% | 55% | 53% | 53% | 50% | 51% | 51% | 51% | 51% | 52% | 51% | 52% | 52% | 73% | 73% |

Table S2: Sequence similarity matrix of Ndk proteins across selected species. Heatmap showing pairwise amino acid identity (%) between Ndk proteins from diverse organisms. *A*: *Aspergillus*, *C. trachomatis*: *Chlamydia trachomatis, C. sequanensis: Criblamydia sequanensis, E. coli*: *Escherichia coli, E. lausannensis*: *Estrella lausannensis, M*: *Mycobacterium*, *P. acanthamoebae*: *Parachlamydia acanthamoebae*, *P. aeruginosa*: *Pseudomonas aeruginosa*, *S*: *Simkania*, *W*: *Waddlia*.
